# A rapid, low-cost approach to solid immersion lens fabrication for enhanced resolution in optical microscopy

**DOI:** 10.64898/2026.02.19.706816

**Authors:** Liam M. Rooney, Jay Christopher, Shannan Foylan, Charlie Butterworth, Laura Copeland, Lewis D. Walker, Katie Coubrough, The SOMC 2025 Consortium, Gwyn W. Gould, Margaret R. Cunningham, Ralf Bauer, Gail McConnell

## Abstract

Solid immersion lenses (SILs) enhance the spatial resolution of an optical microscope by increasing the effective numerical aperture (NA) without physical modification of the objective lens. However, SIL application remains limited by cost, fragility, and accessibility. We present a rapid, single-step fabrication process to create optical quality hemispherical SILs using consumer-grade UV-curable transparent resin which reduces material costs by over five orders of magnitude relative to commercial glass counterparts. Our method produced resin SILs within seconds which can be easily implemented into conventional microscopy setups for increasing the effective NA. Quantitative imaging of USAF resolution targets and histology muscle preparations demonstrated a resolution enhancement approaching theoretical limits and comparable performance to N-BK7 glass SILs. This enabled visualisation of features usually below the diffraction limit of low NA dry objectives at a fraction of the cost of otherwise required high-powered objective lenses. To demonstrate accessibility and translational potential, our workflow was taught in a practical tutorial of an international microscopy course, where non-expert participants successfully fabricated, characterised, and applied SILs within a single session, reporting high confidence in independent implementation. We established ultra-low-cost resin SILs as a practical, scalable option to enhance the spatial resolution of routine optical microscopes and as an accessible and cost-effective platform for optics education.

## Introduction

The resolving power of an optical microscope is limited by the collection angle of the objective lens and the refractive index of the object space, together defining the numerical aperture of the objective lens (NA_*Obj*._). Low NA objectives (typically NA_*Obj*._ < 0.8) are routinely used to image specimens in air (i.e., refractive index, *n*, of object space, *n* = 1.0), whereas higher-NA objective lenses mostly require immersion in water (*n* = 1.33), silicone oil (*n* = 1.40), glycerol (*n* = 1.47), or oil (*n* = 1.51-1.55). These immersion media increase the angular range over which light is collected from the specimen plane into the objective, enabling higher spatial resolution imaging ^1^.

Solid immersion lenses (SILs) offer an alternative approach to liquid immersion strategies. Typically, a hemispherical solid lens is placed directly onto the specimen, increasing the refractive index of object space without modifying the objective lens itself ^2,3^. Resolution enhancement is achieved through the same basis as liquid immersion, whereby light from the specimen is collected over a wider range of angles before entering the objective ^4^. In addition, the SIL increases the total magnification of the imaging system in line with the refractive index of the SIL. Effective implementation of a hemispherical SIL requires three key parameters: the refractive index of the SIL must exceed that of the surrounding medium; the first surface, *R*_1_, must be planar with a hemispherical second surface (radius of curvature, *R*_2_, = radius); and the overall size of the SIL must be compatible with the working distance of the objective lens ^5^. Despite their advantages, SILs are rarely used in routine microscopy due to their fragility, cost, and lack of familiarity among end users. Commercial SILs are typically fabricated from optical-quality N-BK7 glass and cost approximately £50 per lens, presenting a significant barrier for widespread adoption, particularly in teaching laboratories and resource-limited settings. Given recent advances in the use of additive manufacturing for optical-quality lens fabrication ^6–9^, we reasoned that the barrier to entry for SIL imaging could be lowered by using consumer-grade transparent resin and by creating a rapid manufacturing process accessible to non-specialists.

We aimed to create a simple, rapid and low-cost fabrication method that enabled end users to produce and implement resin SILs while retaining the optical performance of commercial glass elements. We present a rapid fabrication process for manufacturing optical-quality SILs from transparent, consumer-grade UV-curable resin. Following comparison of the resin SILs with their glass counterparts, we demonstrated their ability to enhance spatial resolution using both resolution test targets and biological specimens. Finally, we showed the ease of implementation and potential for broad adoption by integrating the fabrication process into a practical workshop delivered as part of the 2025 Strathclyde Optical Microscopy Course (SOMC). Participants from diverse disciplinary backgrounds were able to fabricate, validate and apply resin SILs within a single session, highlighting the accessibility and translational potential of this approach for enhanced biological imaging.

## Methods

### Background Theory for Enhanced Resolution using Resin-based SILs

Solid immersion lenses serve to increase the effective NA (NA_*Eff*._) of the imaging system by introducing a single hemispherical element between the specimen and the objective lens. The higher refractive index medium of the SIL increases the range of angles collected by the objective lens, compared to a given conventional low-NA dry objective ^10^. The SIL does not introduce an additional aperture but, instead, it scales the object-space NA of the objective proportional to the refractive index of the SIL material (*n*_SIL_) ^11^.

For a combination of a dry objective (i.e., with *NA*_*air*_= *n*_*air*_ sin *θ* refractive index of air and *θ* = the half-angle of collection of the objective lens) and a SIL of refractive index, *n*_SIL_, the maximum angle in object space is preserved inside the SIL, but the effective NA in object space (i.e., NA_*Eff*._) becomes

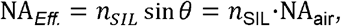

up to the limit defined by the refractive index of the SIL ^10,12^. This follows from Snell’s law: the maximum acceptance angle of the objective remains fixed, but a higher refractive index shortens the in-medium wavelength and increases the accessible spatial frequencies. In practice, this means that even low-to-moderate NA objectives can achieve significantly improved resolving power ^13^ when used with a SIL of suitable refractive index and surface quality.

The effect of the SIL can be described within the context of the Abbe theorem for a diascopic transmission microscope configuration ^14,15^. The minimum resolvable periodicity of a structure (*p*_ffiln_) under a given wavelength of incident light (λ) is given by

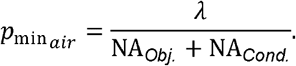

The SIL modifies the objective term by replacing NA_*Obj*_ with NA _*Eff*_., such that

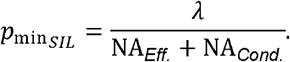

The condenser NA remains unchanged. No additional term is introduced for the SIL because it does not collect diffraction orders independently; it only increases the angular range over which light is collected by the objective.

The resin-based SILs used in this study have refractive indices in the range of common optical glasses (≈ 1.51). For example, a 0.50 NA dry objective used with a resin SIL of *n*_SIL_ = 1.51 results in an effective NA of approximately 0.755. When implemented into a diascopic brightfield transmission microscope aligned for Köhler illumination, this is sufficient to reduce the minimum resolvable period by approximately 20%, such that without SIL (λ = nominally 550 nm):

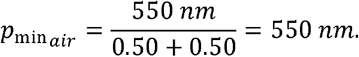

with SIL (λ = nominally 550 nm):

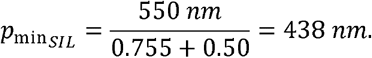

This resolution improvement can be realised without modifying the microscope hardware, provided the SIL is in contact with the coverslip of the sample carrier, the surface curvature is radially symmetrical, and it has reasonable hemispherical tolerance.

### A Rapid Fabrication Process to Produce Resin SILs

A microscope slide was used as the fabrication surface of the resin SILs. The slides were cleaned using neat anhydrous methanol (M/4085/17; FisherScientific, USA) and lens tissue (MC-5; ThorLabs, USA). The maximum height (ergo, volume) of the SIL was dictated by the working distance of the objective lens used for imaging; for the UPLFLN 20×/0.50 NA objective lens (Evident, Germany) used in this study, that was limited to less than 2 mm. The volume of the hemispherical SIL was calculated for a lens with radius of 1.5 mm, requiring 7 μL of resin. The end of a 10 μL pipette tip was cut at a 45° angle using a pair of sharp scissors to minimise pipetting errors of the viscous resin. The 7 μL of Clear UV Resin (4^th^ generation; VidaRosa, China) was deposited onto the clean glass slide and the assembly was rapidly cured for 5 seconds using a UV LED torch (λ = 365 nm, 3 watts) (PR110019-2; LE Innovation Ltd., Ireland), placed approximately 50 mm from the resin surface. The cured SIL was removed from the glass slide by freezing for 2 minutes at -20°C and gently flexing the glass slide so that SIL was released and could then be moved onto the specimen for imaging. Figure 1 shows the principles of using a SIL to enhance the NA_*Eff*._ of the imaging system and a schematic of the rapid manufacturing process.

**Figure 1.**
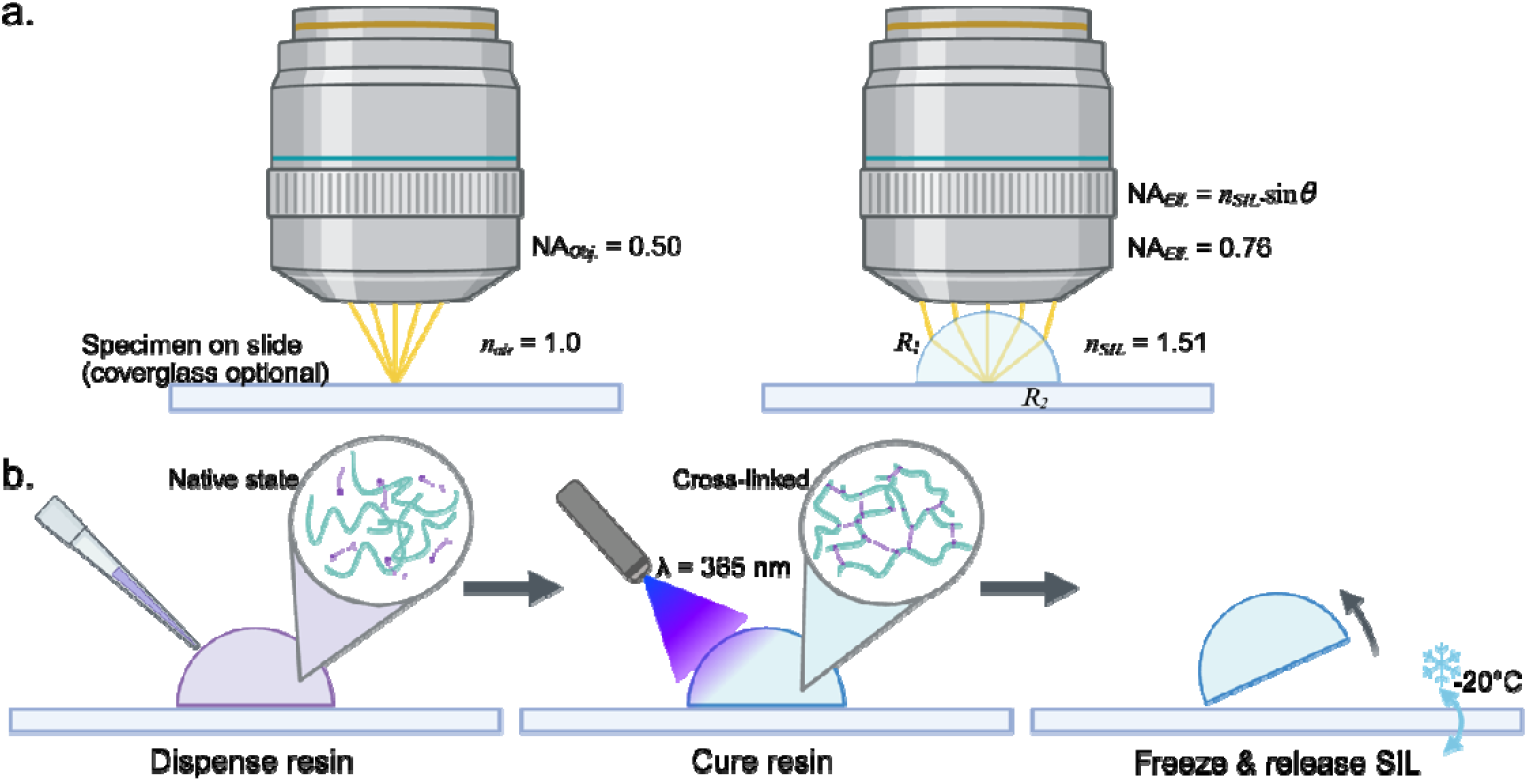
The principles and rapid fabrication of resin SILs. **(a)** A schematic comparing the brightfield microscopy setup for conventional low magnification dry objective lenses, where the resolution is dependent on the half-angle of the collected light through refractive index, *n*_*air*_, and for a resin SIL configuration, where resolution is enhanced owing to the increased effective NA afforded by the increased refractive index of the SIL compared to air. **(b)** A flow diagram describing the rapid manufacturing process for resin SILs: the required volume of transparent resin is deposited onto a clean microscope slide, cured using ultraviolet light and removed by freeze-thawing and flexing the slide.

### Interference Reflection Microscopy to assess surface curvature and quality of resin SILs

Interference Reflection Microscopy (IRM) facilitated high resolution 3D reconstructions of the surface profile of the SIL lenses and comparison of the radius of curvature (*R*_*2*_) versus theory. The principles of IRM are detailed elsewhere ^16,17^. Briefly, a resin SIL was placed convex side down on a Type 1.5 coverglass that spanned the stage insert of an IX81 inverted microscope coupled to an FV1000 confocal laser scanning unit (Olympus, Japan). The reflected light was collected by using an 80/20 beamsplitter in place of the dichroic filter and imaged using a 10×/0.40 NA objective lens. A 458 nm argon laser (GLG3135; Showa Optronics,Japan) was used as the incident light source, which was reflected from refractive index boundaries at the specimen plane. The reflection signal was detected using a PMT with the detection limited to 458 ± 5 nm. Reflected light from different positions along the optical axis exhibited constructive and destructive interference depending on the optical path difference between the reflected beam from the SIL surface and the underlying coverglass. The resulting interference fringes created a 2D projection of the 3D topography of the SIL surface, where interference orders were separated along the optical axis. The axial separation of constructive and destructive interference orders is described below, respectively, where *z* = order spacing, *N* = order, λ = wavelength of incident light, and *n*_*m*_ = refractive index of the imaging medium (in this case, air = 1.0).

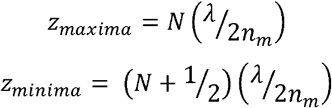

### Comparing the optical performance of glass and resin SILs

A N-BK7 glass hemisphere with a diameter of 3 mm (47-269; Edmund Optics, USA) was used to compare the performance of the resin SILs. Both glass and resin SILs had the same volume, diameter and refractive index. A USAF resolution test target (15b; ReadyOptics, USA) was used to determine the resolution enhancement of both glass and resin SILs when used with a routine low magnification brightfield transmission microscope. A BX60 upright brightfield microscope (Olympus, Japan) was aligned for Köhler illumination, such that the NA_*Cond*._ was equal to NA_*Obj*._. Brightfield transmission images were acquired using a 20×/0.50 NA objective lens (Olympus, Japan) via a monochrome CMOS camera (UI-3060CP-M-GL R2; IDS Imaging, Germany). Images of the USAF target slide were acquired in air, with a glass SIL and with a resin SIL. The resolution enhancement facilitated by the SILs was determined by measuring the peak prominence of the USAF gratings, measuring at which point the elements within the test groups become indistinguishable .

The performance of the resin SIL was further compared by imaging a thin section preparation of toluidine blue-stained mouse muscle tissue mounted in Histomount (Invitrogen, USA). The specimen was chosen specifically because of the fine striated structures that are indiscernible in conventional low-magnification imaging. The specimen was imaged using the same brightfield transmission setup as described above, except with a colour CMOS camera (CS165CU; ThorLabs, Germany) to visualise the histological staining. Images were acquired of the specimen in air and with glass or resin SILs placed directly on top of the coverglass. The muscle striations were detected by generating an intensity plot profile over the length of a section of muscle tissue and comparing the same region between air imaging and SIL imaging. Intensity plot profiles were generated using FIJI^18^ (v2.16.0).

### Implementing and evaluating rapid SIL manufacturing in an educational setting

To demonstrate the ease of implementation and the value of rapid SIL manufacturing as an educational resource for optical engineering, we designed a short practical workshop as part of the 2025 Strathclyde Optical Microscopy Course. Twenty-four students from various backgrounds and career stages (spanning physics, biology, chemistry and engineer; from postgraduate students to senior microscopy facility managers) were provided a survey before and after the workshop (See Supplementary Information). This was designed to assess participants’ knowledge of optical engineering, SIL imaging, and 3D printing. We provided a brief tutorial covering additive manufacturing, the benefits of SIL imaging, and how to fabricate SILs using the protocols described above. Students were provided with aliquots of clear resin, clean microscope slides, pipettes and pipette tips, a UV LED torch, and a 3D printed opaque shielding box to prevent UV exposure during curing. The students were given 45 minutes to calculate the volume required according to the prescription needed for a given brightfield transmission microscope. They were then provided with a simple optical setup of a collimated laser source and asked to measure the focal length of their SIL lens and verify that it matched the theoretical value from the desired lens prescription. Students were asked to complete a post-workshop survey to capture what they had learned, how accessible they found it, and if it had altered their perception of additive manufacturing, SIL imaging or optical engineering for their own research projects. Qualitative assessment of the survey responses focussed on extracting benefits of the workshop and areas of improvement. Quantitative analysis of the survey responses focused on rating the students’ understanding of core concepts after the workshop and measuring the ease of manufacturing that would enable researchers to replicate and implement SIL fabrication in their own imaging research.

## Results

### Resin SIL Performance is Comparable to Glass SILs

A USAF Resolution Test Target was used as a comparative performance test for resin and glass SILs. Figure 2 shows a comparison of the test specimen imaged in air using a standard 20×/0.5 NA objective lens (Figure 2a), with a glass SIL (Figure 2b) and with a resin SIL (Figure 2c). The resolution was measured by assessing a line intensity plot profile through the USAF group(s) containing the theoretical resolution limit (Group 9, Element 5; 615 nm line spacing – diffraction limit for 0.5 NA objective lens; Group 10, Element 1; 488 nm line spacing – theoretical resolution limit for the effective NA of this SIL setup) (Figure 2d-e). Intensity plot measurements showed that both glass and resin SILs enhanced the resolution of the imaging system, up to the theoretical improvement limit. The data show that the improvement was more evident in the commercial glass SIL, as evidenced by reduced contrast in USAF groupings, but the resolution enhancement was also demonstrable with the resin SIL. This demonstrates that a single low-cost rapid-fabrication lens can achieve almost comparable resolution enhancement to more costly and restrictive glass elements. Homogeneity of illumination was retained across all imaging conditions (>95% homogeneity across the field of view) and IRM confirmed the surface curvature of the resin SIL matched the expected theoretical curvature (Supplementary Figure 1).

**Figure 2.**
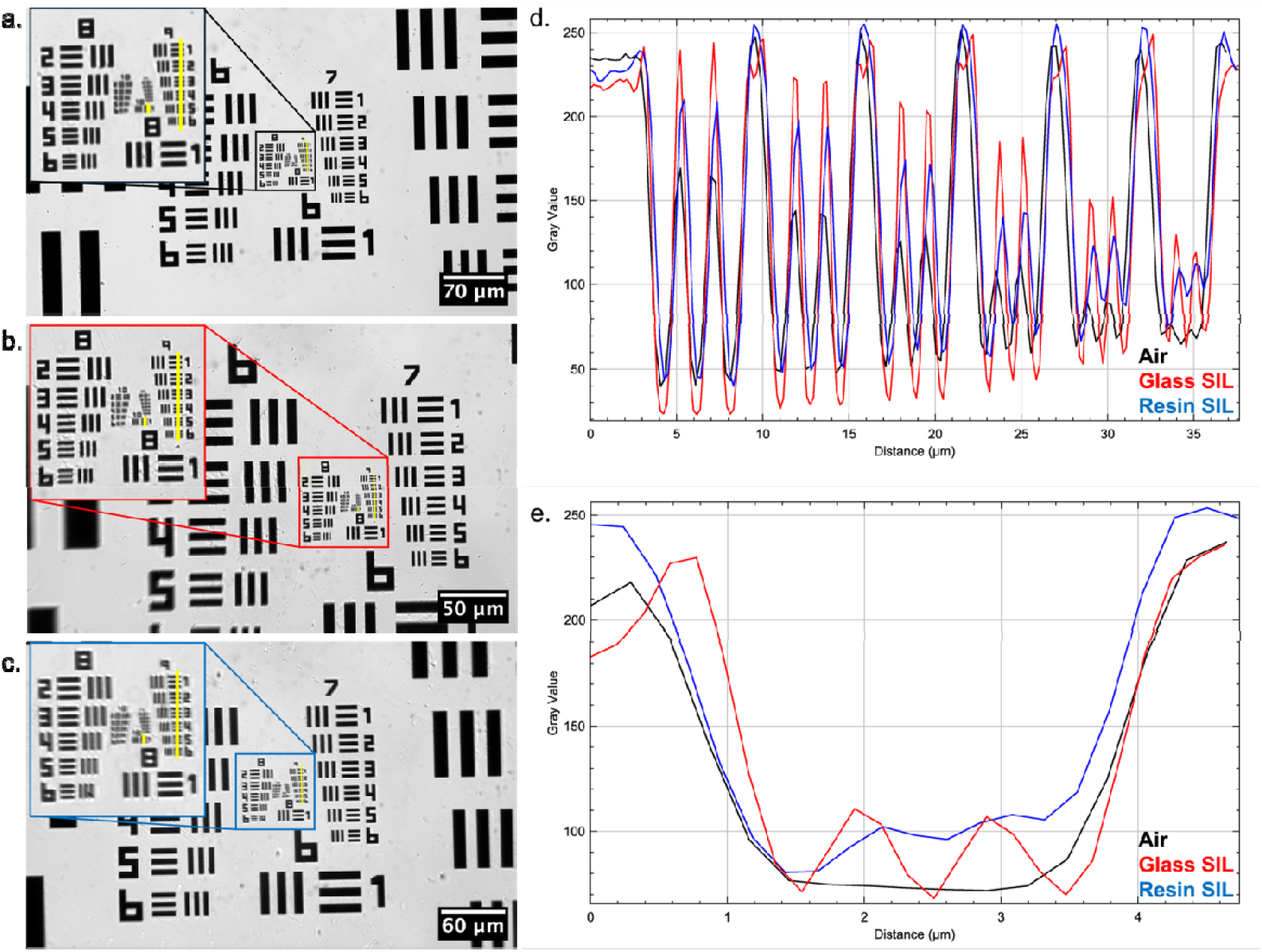
Comparison of the imaging performance of resin and glass SILs. A USAF 2015a Resolution Test Target imaged using a standard brightfield transmission setup with a 20×/0.5 NA objective lens in **(a)** air, **(b)** with a glass SIL, and **(c)** with a resin SIL. Digitally magnified regions of interest are presented that show measurement traces (yellow) for intensity plot profiles. An intensity plot profile for **(d)** Group 9 and **(e)** Group 10, Element 1 are shown, with traces shown for air (black), glass SIL (red) and resin SIL (blue). Line thickness = 5 pixels.

### Low-cost Resin SILs Facilitate Visualisation of Sub-Diffraction Limit Structures

The resolution enhancement and ease of implementation was demonstrated using a thin section histology preparation of mouse paw skeletal muscle. The arrangement of the contractile machinery of the muscle tissue, the sarcomere, forms periodic banding structures with approximately 1 μm intervals along the long axis of the muscle cells, which are difficult to detect using low-NA dry objective lenses (Figure 3a). However, the use of a resin SIL enhanced the spatial resolution such that the repeating banded structure was detected throughout the muscle preparation with no additional changes to the microscope (Figure 3b). The performance of the low-cost rapid-fabrication resin SIL was comparable to that of the glass SIL (Supplementary Figure 2).

**Figure 3.**
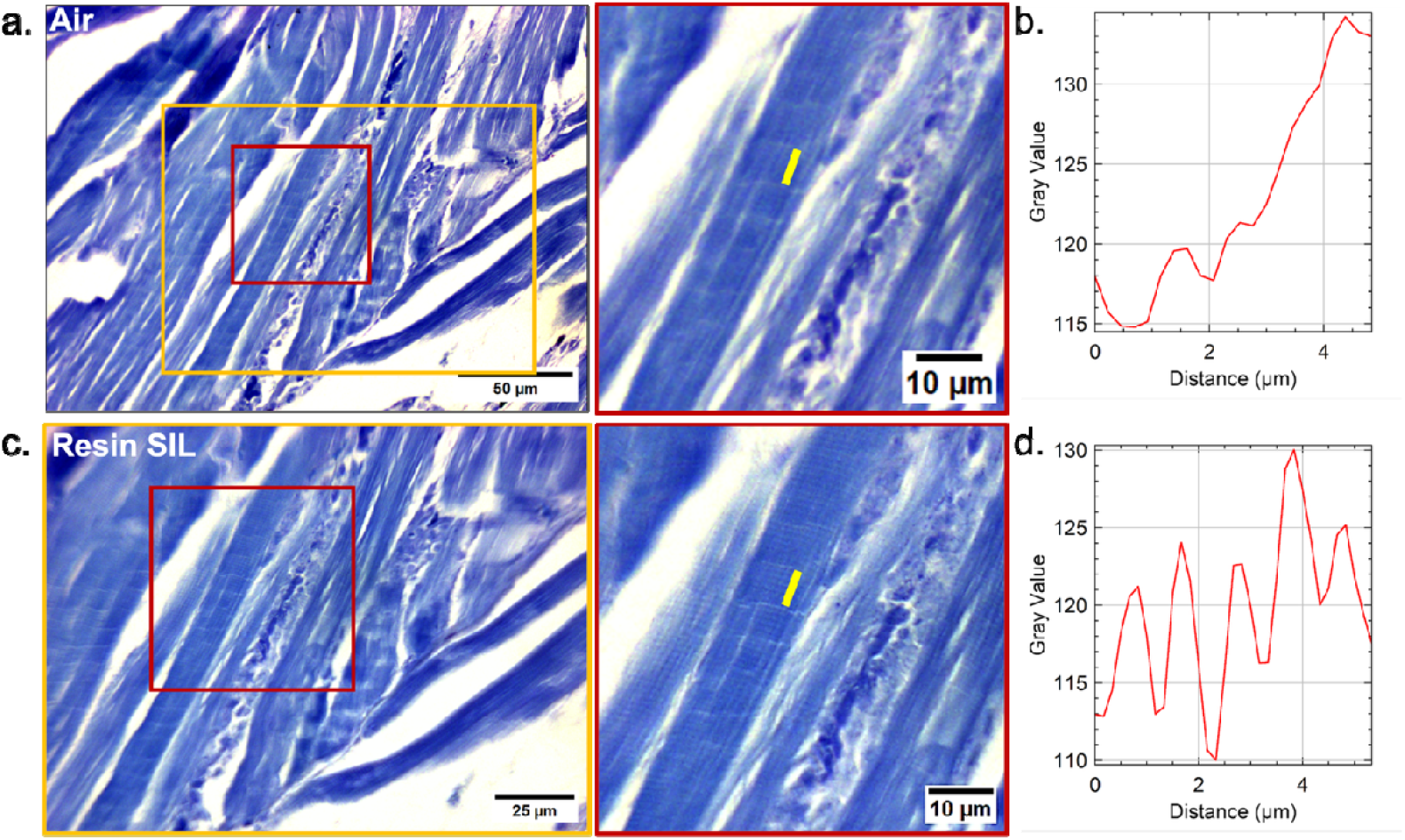
Low-cost, rapid-fabrication SILs facilitate detection of sub-diffraction limited biological structures. A mouse paw muscle histology preparation was imaged in **(a)** air using a 20×/0.5 NA objective lens. A digitally magnified region of interest from the image is designated by the area in red, with a yellow trace showing the region from which the intensity along the long axis of the muscle cell was measured. **(b)** A plot of the intensity profile marked in (a) reveals no detected sarcomere banding structures along the axis of the muscle cell. **(c)** The same region was imaged using a resin SIL, which slightly magnified the field of view (shown compared to the original field of view by the yellow box in (a)). The same digitally magnified region and measurement area were chosen to provide a comparison between conventional dry imaging and SIL imaging. **(d)** A plot of the intensity profile marked in (c) shows the previously unresolved repeating banded structure of the sarcomeres along the muscle fibres.

### Resin SIL Production is Accessible and Easily Implemented

To demonstrate the ease of the manufacturing method and its implementation, we designed a practical workshop and embedded it into the framework of an existing international microscopy course, the Strathclyde Optical Microscopy Course. The cohort had little-to-no prior experience of additive manufacturing or lens fabrication, with approximately 50% of the participants being life scientists with no prior knowledge of optics. We collected pre- and post-workshop surveys from the students which contained qualitative and quantitative assessment of their understanding, ability to replicate the manufacturing process, and confidence of implementing resin SILs in their own institutes after the course. Figure 4 shows the change in understanding of lens fabrication and confidence to implement SILs from the cohort before and after the workshop. We showed that 95% of students had an increase in their understanding of the fabrication process, with 90% of the cohort scoring 3 and above (out of 5) in their understanding. All students reported that they felt some degree of confidence in replicating the SIL fabrication and implementing it after the course, with 75% scoring themselves as ‘confident’ or ‘very confident’. Participants from diverse backgrounds successfully fabricated and applied SILs during the session, highlighting the practical accessibility and reproducibility of the workflow.

**Figure 4.**
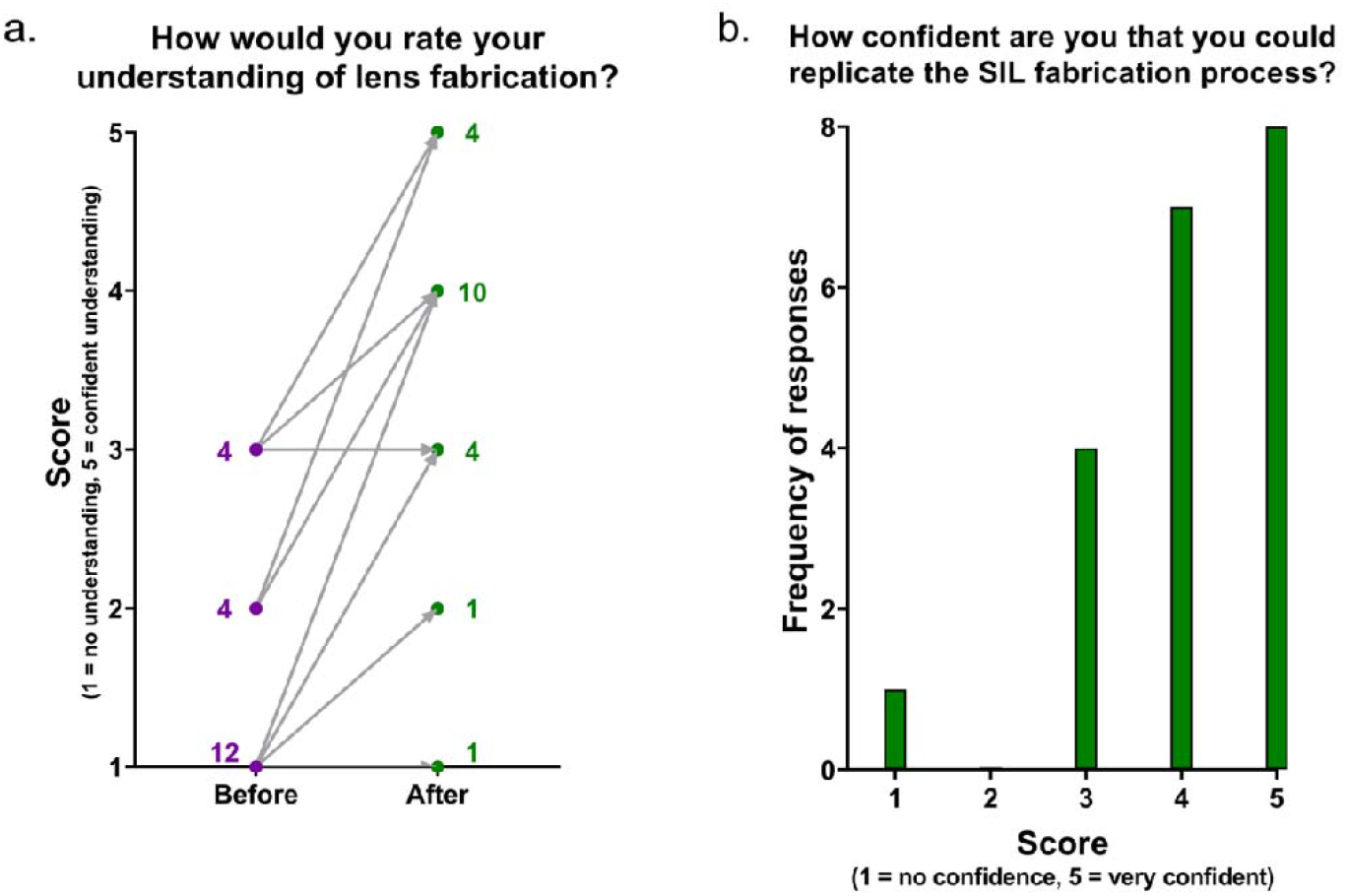
Quantitative assessment of student learning and implementation of resin SILs. **(a)** A self-scored conversion plot showing increased understanding of the cohort before and after the SIL fabrication workshop. **(b)** A score of the confidence of students to replicate SIL fabrication and implementation following completion of the fabrication workshop.

Qualitative free-text responses from the pre- and post-workshop surveys revealed a clear shift in the perceptions of participants for both solid immersion lenses and additive manufacturing for microscopy. Prior to the workshop, respondents frequently expressed uncertainty around optical quality, surface precision, and robustness of self-fabricated components, alongside a general perception that lens manufacturing and optical design were specialist activities. Post-workshop responses instead emphasised the simplicity, speed, and accessibility of the single-step resin SIL fabrication process, with many participants highlighting surprise at how quickly functional optical elements could be produced and deployed. A recurring theme was the perceived feasibility of integrating resin SIL fabrication into existing laboratory workflows, particularly for low-cost resolution testing, teaching, outreach, and rapid prototyping. Participants from life sciences backgrounds noted increased confidence in engaging with optical concepts such as NA and resolution, while those with prior engineering or physics experience highlighted the value of the method as a flexible and low-risk route for experimentation. Across disciplinary backgrounds, respondents consistently framed the approach as enabling and democratising, reinforcing the potential for broad adoption of SILs in both research and educational settings.

These data demonstrate that resin SIL manufacturing is accessible, reproducible and implementable across a variety of skill levels and disciplinary backgrounds, evidencing great potential for wide uptake and implementation of resin SILs to improve the imaging performance of routine microscope setups.

## Discussion

Solid immersion lenses offer the potential to enhance the spatial resolution of low-NA objective lenses with minimal change to the optical setup, but they are seldom used due to their expense, fragility and perceived implementation barriers. We demonstrated a rapid, single-step fabrication process to create ultra-low-cost resin SILs which facilitated comparable resolution improvements to commercial glass alternatives for routine brightfield transmission microscopy. Implementation of the resin SIL effectively increased the NA of a low-NA objective lens by 20%, resolving biological structures otherwise undetected by conventional imaging with a dry configuration. The measured resolution improvement conformed with the theoretical enhancement based on scaling the effective NA by altering the refractive index in object space, confirming that consumer-grade UV-curable resins possess suitable optical properties for SIL-based resolution enhancement. Measurements on USAF resolution targets showed that both resin and glass SILs enabled detection of spatial structures approaching that were otherwise unresolved using a conventional air objective setup. Moreover, we demonstrated the ease of uptake and implementation by introducing a practical workshop for a cohort of mixed discipline students. All students reported that they were confident with their ability to produce and use resin SILs and 95% demonstrated an improvement in their understanding of SIL manufacturing following completion of the workshop.

A defining outcome of this study is the dramatic reduction in cost achieved by the resin-based fabrication approach. At the time of writing, commercially available N-BK7 hemispherical SILs are priced at approximately £50 per lens. In contrast, the resin SILs described here require approximately 7 µL of material, resulting in material costs of ∼£0.00020 per lens (VidaRosa resin = £30/kg). Even when accounting for ancillary fabrication materials (totalling approximately £5.00), including microscope slides, solvents, lens tissue and a consumer-grade UV curing torch, this represents a real-terms cost reduction of over five orders of magnitude. This cost reduction fundamentally alters the accessibility of SIL imaging, enabling disposable lenses, rapid iteration, and application-specific fabrication without the financial or practical risks associated with fragile glass elements.

The optical performance of the resin SILs was slightly lower than that of commercial glass lenses at the highest spatial frequencies, as evidenced by reduced modulation contrast in the finest USAF groupings. Interference Reflection Microscopy revealed that the single step curing process reliably produced lenses with curvature profiles close to the intended hemispherical geometry, though small deviations in radial symmetry were detected. These deviations are likely to introduce some non-spherical aberrations or surface-induced phase errors, which become increasingly significant at higher NAs ^19^. Such aberrations are not inherent limitations of resin materials and may be mitigated through improved control of droplet volume, curing dynamics, or surface energy during fabrication. In future implementations, adaptive optics or computational wavefront correction strategies could further compensate for residual aberrations ^20,21^, particularly in applications requiring larger fields of view or quantitative intensity measurements.

The refractive index of the resin used in this study (≈ 1.51) is comparable to that of standard optical glasses, enabling substantial NA enhancement when paired with dry objectives. Higher-index SIL materials have been reported, including metal oxide and glass substrates that enabled effective NAs far exceeding the design limitations of standard objective lens specifications ^11^. Examples of high-index materials currently used in SIL manufacturing include sapphire (monocrystalline α - aluminium oxide) ^4,22,23^, cubic zirconia (zirconium dioxide) ^24^, silicon ^25–28^, and titanium dioxide ^29,30^, with applications spanning the visible, near infrared and terahertz ranges. While these systems can provide further resolution gains, they typically require specialised materials, precision machining, or complex integration workflows that limit their use to specialist laboratories. In contrast, resin-based SILs prioritise ease of fabrication, safety, and compatibility with existing microscopes, offering a favourable balance between achievable resolution enhancement and practical usability.

Alternative SIL geometries, such as super-hemispherical (Weierstrass) designs ^4,10^, aspherical lenses ^10^ and *in situ* fabricated SILs produced via moulding or lithographic methods ^13^ have shown improved aberration control, extended NA scaling, or increased magnification. However, these approaches often require multi-step fabrication processes, cleanroom-compatible techniques, or specialised instrumentation. The single-step droplet-based approach presented here provides a democratised solution, enabling rapid fabrication and immediate deployment without modification of microscope hardware.

Beyond brightfield transmission microscopy, SILs have been successfully applied to a wide range of imaging modalities, including Raman thermography ^13^, cryogenic imaging ^31^, semiconductor wafer inspection ^24^, and optical data storage ^32,33^. SIL-enhanced fluorescence ^34–37^ imaging has enabled improved signal collection and resolution in systems where high-NA immersion objectives are impractical. Additional SIL applications have seen use as micro-optics ^38^, in generating sub-micron structured illumination patterns ^39^ and in terahertz imaging ^40^. The compatibility of resin SILs with such modalities represents an important direction for future work, particularly in low-cost, portable, or field-deployable imaging systems where conventional immersion optics are unavailable.

The successful integration of resin SIL fabrication into an educational workshop further highlights the broader impact of this approach. Participants from diverse disciplinary backgrounds, many with no prior experience in optics or lens manufacturing, were able to fabricate, characterise, and apply SILs within a single practical session. The significant increase in confidence and understanding reported in post-workshop surveys demonstrates that the method functions not only as a research tool but also as an effective educational platform for introducing core optical concepts such as numerical aperture, refractive index, and diffraction-limited resolution. This accessibility supports wider dissemination of advanced microscopy techniques and encourages adoption across disciplines.

Future developments may focus on improving fabrication reproducibility through controlled dispensing, exploring alternative resin formulations with higher refractive indices, and extending the method to more complex SIL geometries. Combining resin SILs with adaptive optics, computational correction, or structured illumination methods may further expand their applicability and performance to further democratised other high-resolution modalities. Together, these developments position ultra-low-cost resin SILs as a flexible, scalable, and widely deployable strategy for enhancing the performance of routine optical microscopy systems.

## Acknowledgements

The Authors wish to thank the funders who supported this work. LMR and GM were funded by the Leverhulme Trust. LMR was funded by the University of Glasgow. JC, CB and RB were funded by Biotechnology and Biological Sciences Research Council (BBSRC) (BB/Z51486X/1), an Engineering and Physical Sciences Research Council studentship (EP/T517938/1) and supported by the National Manufacturing Institute Scotland (NMIS). SF, GWG and GM were supported by the BBSRC (BBX005178/1). LC and LDW were funded by an EPSRC iCASE studentship (EP/Y528833/1). KC was funded by the University of Strathclyde. GM was supported by the Medical Research Council (MR/K015583/1) and the BBSRC (BB/P02565X/1 and BBT011602).

The authors would like to thank the SOMC 2025 Student Consortium, who are included as a group author on this manuscript. Their enthusiasm and eagerness to learn enabled us to test our manufacturing process at scale within the framework of the Strathclyde Optical Microscopy course. Thank you to all of you; Emily Horsburgh, Amrutha Sankar, Wenjing Huang, Judy Bagi, Peter Gordon, Abbey Began, Kay Polland, Megan Greer, Enya Berrevoets, Sarah Lecinski, Sarah Gosling, Ewan Drever-Smith, Katherine Baxter, Connor MacDonald, Alistair Ozzie Gemmell, Joseph Navaratne, Nathanael Tan, Juliana Kim, Amy Bottomley, Vincenzo Infante, Charlie Butterworth, Philip Graemer, Naveed Ul Islam, Siân Culley, Beatrice Bottura, and Katherine Paine.

Figure 1 was created in BioRender. Rooney, L. (2026) https://BioRender.com/2ftzqxg.

## Data Availability Statement

The data that support the findings of this study are openly available at the University of Strathclyde KnowledgeBase: doi.org/10.15129/acf9d15e-3b51-4001-ad78-a81c262f77a3

