## Supplementary Information for "A rapid, low-cost approach to solid immersion lens fabrication for enhanced resolution in optical microscopy"

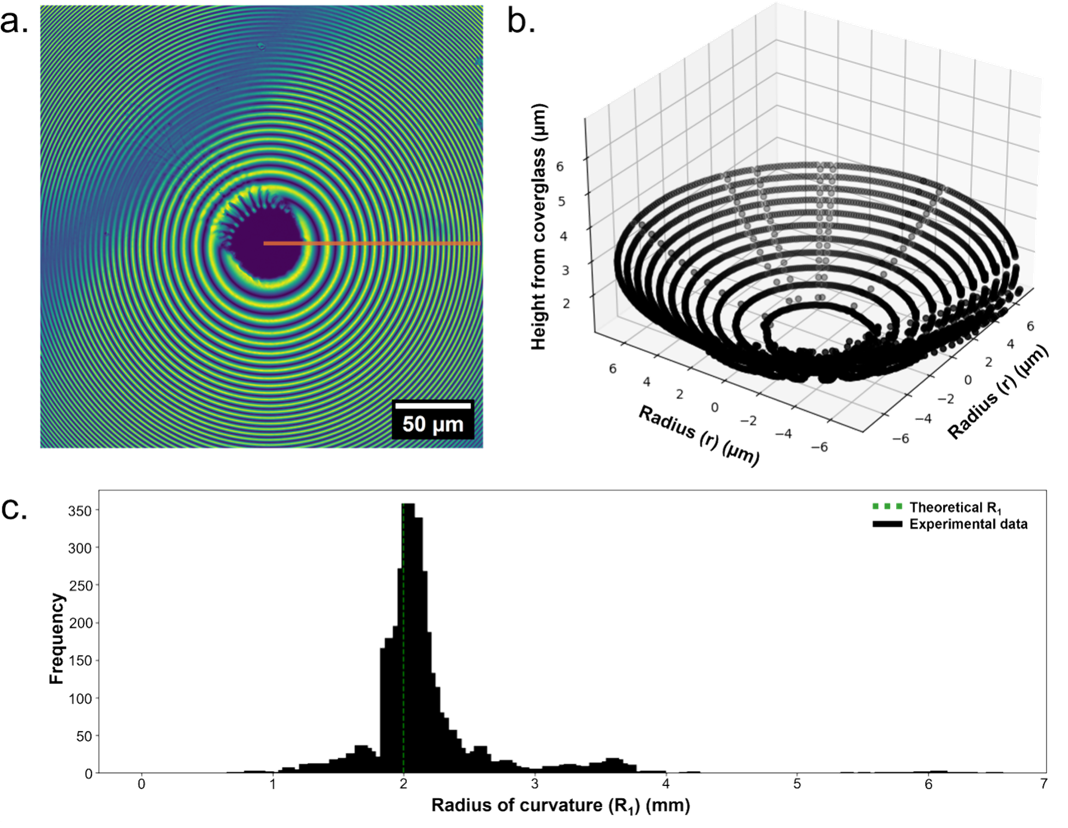

**Supplementary Figure 1. Surface profilometry of a resin SIL using Interference Reflection Microscopy.** **(a)** A false coloured IRM image of a resin SIL surface (λ = 458 nm). Orange ROI designates a given radius used to measure the intensity profile around the circumference of the lens surface. **(b)** A 3D reconstruction of the resin SIL surface computed from the raw IRM image. **(c)** A histogram of the distribution of radius of curvature (*R_2_*) values measured around the circumference of the lens (black) compared to the theoretical *R_2_* (green).

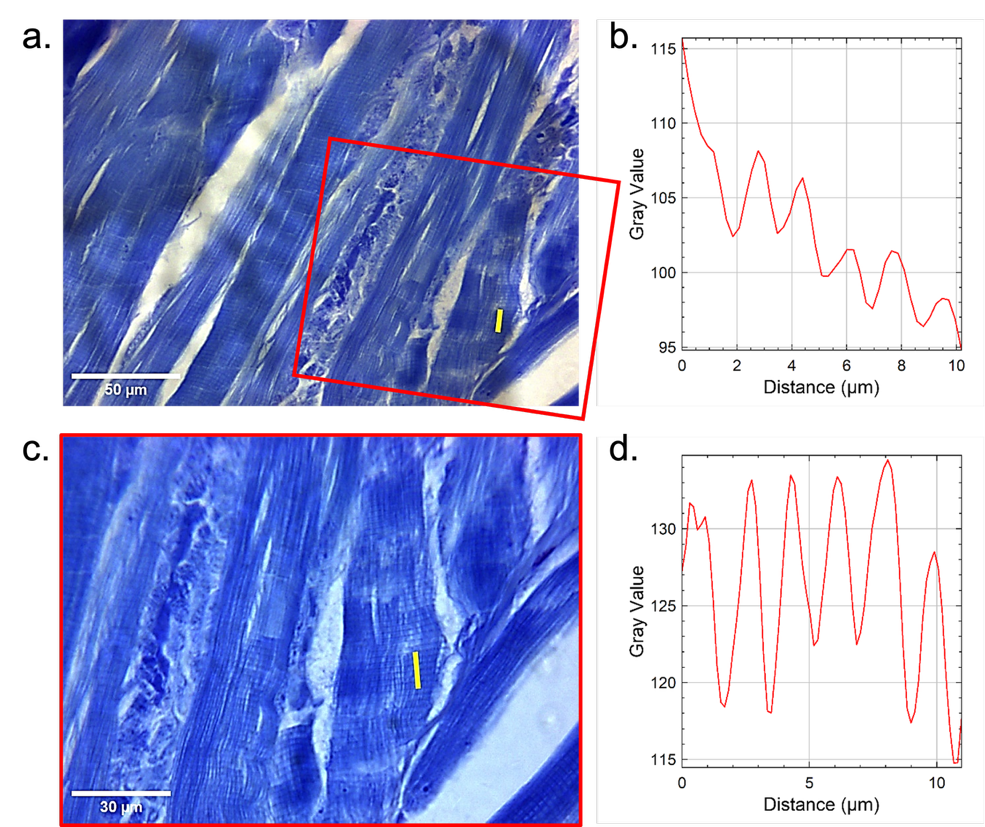

**Supplementary Figure 2. Detection of sarcomere structures using a glass SIL.** A mouse paw muscle histology preparation was imaged in **(a)** air using a 20×/0.5 NA objective lens. A yellow trace showing the region from which the intensity along the long axis of the muscle cell was measured. **(b)** A plot of the intensity profile marked in (a) reveals faint sarcomere banding structures along the axis of the muscle cell. **(c)** The same region was imaged using a glass SIL, which slightly magnified the field of view (shown compared to the original field of view by the red box). The same measurement area was chosen to provide a comparison between conventional dry imaging and glass SIL imaging. **(d)** A plot of the intensity profile marked in (c) shows the previously poorly repeating banded structure of the sarcomeres along the muscle fibres with much improved precision and contrast.

**SOMC survey questions:**

1. **PRE-WORKSHOP**

- How would you rate your current level of experience with 3D printing: None, Basic, Intermediate, Advanced
- Have you previously used 3D printing for microscopy components

If yes, please describe:

- which of these platforms are you familiar with: OpenFlexure, UC2, EnderScope, None of the above, Other
- How would you rate your understanding of 3D printed lens fabrication? (1 (not at all) - 5 (expert))
- Have you heard of, or have you used solid immersion lenses before: yes/no

If yes, in what context

- What do you hope to gain from this workshop (tick all that apply): Practical skills in 3D printing, Insights into open microscopy platforms, understanding lens fabrication, ideas for my own research, other
- How relevant is 3D printed hardware for your research: 1 (not at all) - 5 (a central focus/technique))
- What are you most curious about in the 3D printing workshop session?
- Any concerns about using 3d printed components in microscopy?
- Your disciplinary background: Microbiology, Cell biology, Engineering, Optics, Chemistry, Computer science/image analysis, other

2. **POST-WORKSHOP**

- How would you now rate your understanding of 3D printed lens fabrication? (1 (no understanding - 5 (very confident)
- What did you find most useful or interesting?
- Did the workshop change your perspective on any of the following? Feasibility of printing microscopy components, value of Open Microscopy, Use of 3D printed lenses in research, integration of lens fabrication in lab workflows, other
- Which parts of the session were least clear or could be improved?
- Would you consider using 3D printed optical components in your own research? Definitely not, possibly, likely, definitely

Please briefly explain:

- How confident are you that you cold replicate the single-step solid immersion lens fabrication process in your own lab? (1 (not confident at all) - 5 (very confident)
- Are there any key concepts or ideas that you expect you could apply in your own research?
- How engaging did you find the workshop format and demonstrations (1 not engaging - 5 very engaging
- Was the pacing and amount of content appropriate: too slow, just right, too fast
- Are there any areas of 3D printing/open hardware/open microscopy that you would like to have heard about that we missed? Please explain briefly:
- I consent to the use of my anonymous responses in aggregated form for scientific research purposes, including publication

Full responses available via Strathclyde KnowledgeBase: doi.org/10.15129/acf9d15e-3b51-4001-ad78-a81c262f77a3
